# Robust quantification of multiplexed fluorescent protein-based biosensors in plant tissues

**DOI:** 10.64898/2026.03.13.711581

**Authors:** Valentina Levak, Anže Županič, Karmen Pogačar, Nastja Marondini, Katja Stare, Tina Arnšek, Katja Fink, Kristina Gruden, Tjaša Lukan

## Abstract

Genetically encoded biosensors are one of the essential tools in biological research. They enable visualization of molecules of interest from the subcellular level to entire organism level *in vivo* and can be used to monitor presence of small molecules, gene expression, protein activity, and protein degradation. However, multiplexing fluorescent biosensors in plants is notoriously difficult due to signal bleed-through and strong autofluorescence from chlorophyll. In this study, we investigated the potential of multiplexing biosensors based on the selection of reporter fluorescent proteins. We characterized the emission spectra, fluorescence lifetimes, and relative brightness of diverse fluorescent proteins in plant leaves. We show that selected proteins exhibit comparable brightness, supporting their use in co-expression experiments and reliable quantification of individual signals. To separate three overlapping signals, we applied two different linear unmixing approaches and compared them to results obtained without unmixing. We identified channel separation unmixing approach as the most suitable for biosensors. Additionally, we show how unmixing with the selected approach can be applied to separate autofluorescence and five fluorescent proteins. We further validated this approach in virus-infected cells by following organelle dynamics *in vivo*. Finally, we demonstrate the feasibility of high-throughput segmentation and quantification with a custom MATLAB workflow for nuclei, chloroplasts, and cytoplasm signal analysis. Overall, our work demonstrates that biosensors can be multiplexed, even when their emission spectra overlap.

**Significance statement:** Multiplexing genetically encoded biosensors in plants has been limited by overlapping fluorescent signals and strong autofluorescence. This study presents an optimized framework for linear unmixing and provides a MATLAB-based organelle segmentation tool, allowing precise quantification of multiple fluorescent reporters *in vivo* and advancing real-time visualization of complex cellular processes in plants.

## Introduction

Since its discovery in 1961, green fluorescent protein (GFP) and its homologs from other organisms have become indispensable tools in biological research (Shimomura, 2005). Over the years, the fluorescent protein palette has expanded to include variants spanning from UV-excitable to IR-emitting (Day & Davidson, 2009; Matlashov et al., 2020; Shu et al., 2009). These proteins are widely employed in the development of genetically encoded biosensors, enabling studies with high temporal (millisecond) and spatial resolution across scales—from subcellular to organism level. Despite the availability of a broad range of biosensors, their multiplexed use remains relatively uncommon, particularly in plant research. One of the main challenges arises from the broader excitation and emission spectra of fluorescent proteins compared to spectra of chemical dyes, which complicates simultaneous detection due to spectral overlap. Simultaneous detection of multiple fluorescent proteins in plant tissues—without spectral overlap between the fluorescent proteins or with autofluorescence—is typically limited to two or three fluorescent proteins, each excited by a different laser. However, multiplexing of existing spectrally overlapping biosensors would give an insight into precise spatiotemporal coordination of multiple interacting proteins or processes. Moreover, this approach is particularly advantageous in experimental settings that necessitate the simultaneous use of a large number of fluorescent proteins, for which a sufficient set of variants with non-overlapping spectral properties is not available.

Plant responses to environmental conditions, pathogens, and pests involve a wide range of molecular players, including Ca²⁺, reactive oxygen species (ROS), hormones, and protein kinases (Zhang et al., 2020, 2021). These signalling components interact through sequential cascades and complex crosstalk to regulate and balance diverse physiological processes (Lukan & Coll, 2022). To truly understand plant immune response, we need a way to simultaneously visualize several molecular responses in the same cells at the same time. Multiplexing of genetically encoded biosensors within the same organism and the evaluation of their spatial and temporal interdependence is crucial for understanding how plants integrate multiple signalling pathways (Xiao et al., 2025). The outcome of plant responses to environmental stressors depends not only on the qualitative presence of specific molecules but also on their abundance (Delplace et al., 2020; Jones et al., 2024), making accurate quantification of these biomolecules using genetically encoded biosensors essential. In our observation, the relative brightness of fluorescent proteins strongly influences the successful multiplexing, as sensors with markedly different brightness exhibit unequal sensitivity range of quantification under the same imaging conditions. Moreover, imaging in plant tissues presents additional difficulties due to strong autofluorescence, primarily from chlorophyll and secondary metabolites (Donaldson, 2020).

In this study, we address the challenges of multiplexing genetically encoded biosensors based on fluorescent proteins in plants and suggest optimal protocol for imaging considering the error associated with it. We characterized eight fluorescent proteins—mTagBFP2, mTurquoise2, Venus, mKO2, mCherry, mKate2, mCardinal, and miRFP713—in *Nicotiana benthamiana* leaves, evaluating their emission spectra, fluorescence lifetimes, and brightness. From this set, Venus, mKO2, and mKate2 were also assessed in *Solanum tuberosum* in addition to *N. benthamiana*. To separate overlapping signals, we successfully applied two different linear unmixing approaches. Our protocols could separate overlapping fluorescent protein signals as well as fluorescent protein signal and autofluorescence. Finally, we demonstrate the feasibility of high-throughput segmentation and quantification with a MATLAB workflow for nuclei, chloroplasts, and cytoplasm signal analysis.

## Results and Discussion

### The emission spectra of fluorescent proteins depend more on the imaging system than on the plant matrix

Efficient separation of fluorescent proteins depends on differences in their excitation and emission spectra or fluorescence lifetimes (Tau, τ), as well as comparable brightness to ensure accurate spectral unmixing. Therefore, we first checked if the published emission spectra are comparable to the ones acquired with different confocal systems and in different plant matrices. Additionally, we assayed the fluorescent proteins and autofluorescence of chlorophyll for their fluorescence lifetimes to provide experimental evidence that they can be separated based on their lifetimes *in planta*.

We selected a palette of reported highly bright proteins (mTagBFP2, mTurquoise2, Venus, mKO2, mCherry, mKate2, mCardinal, and miRFP713) and measured their emission spectra and fluorescence lifetimes in plant leaves to detect any potential matrix dependent effects. To enable quantitative imaging, each protein was localized in *N. benthamiana* nuclei using N7 nuclear localization signal and expressed under strong constitutive CaMV 35S promoter in a transient setup. Imaging was performed using Leica’s TCS LSI, Stellaris 5, and Stellaris 8 systems. All proteins except miRFP713 were detectable by all three imaging systems under these conditions; the lack of miRFP713 signal likely reflects insufficient cytoplasmic and nuclear biliverdin, its required cofactor (Matlashov et al., 2020). Notably, we successfully detected mCardinal in plant leaves for the first time, while mKO2 and mTagBFP2 detection had been previously reported *in planta* (Rigoulot et al., 2021). The use of other fluorescent proteins is well-documented in plants (Brunoud et al., 2012; Ghareeb et al., 2016; Mylle et al., 2013).

We first compared the emission spectra of the selected fluorescent proteins obtained by different confocal systems. Consistent with previous observations (Mylle et al., 2013), we noted discrepancies relative to the spectra reported in FPbase (Lambert, 2019), particularly in the red range when using HyD S detectors (Figure 1a, Stellaris 5 and Stellaris 8).

**Figure 1:**
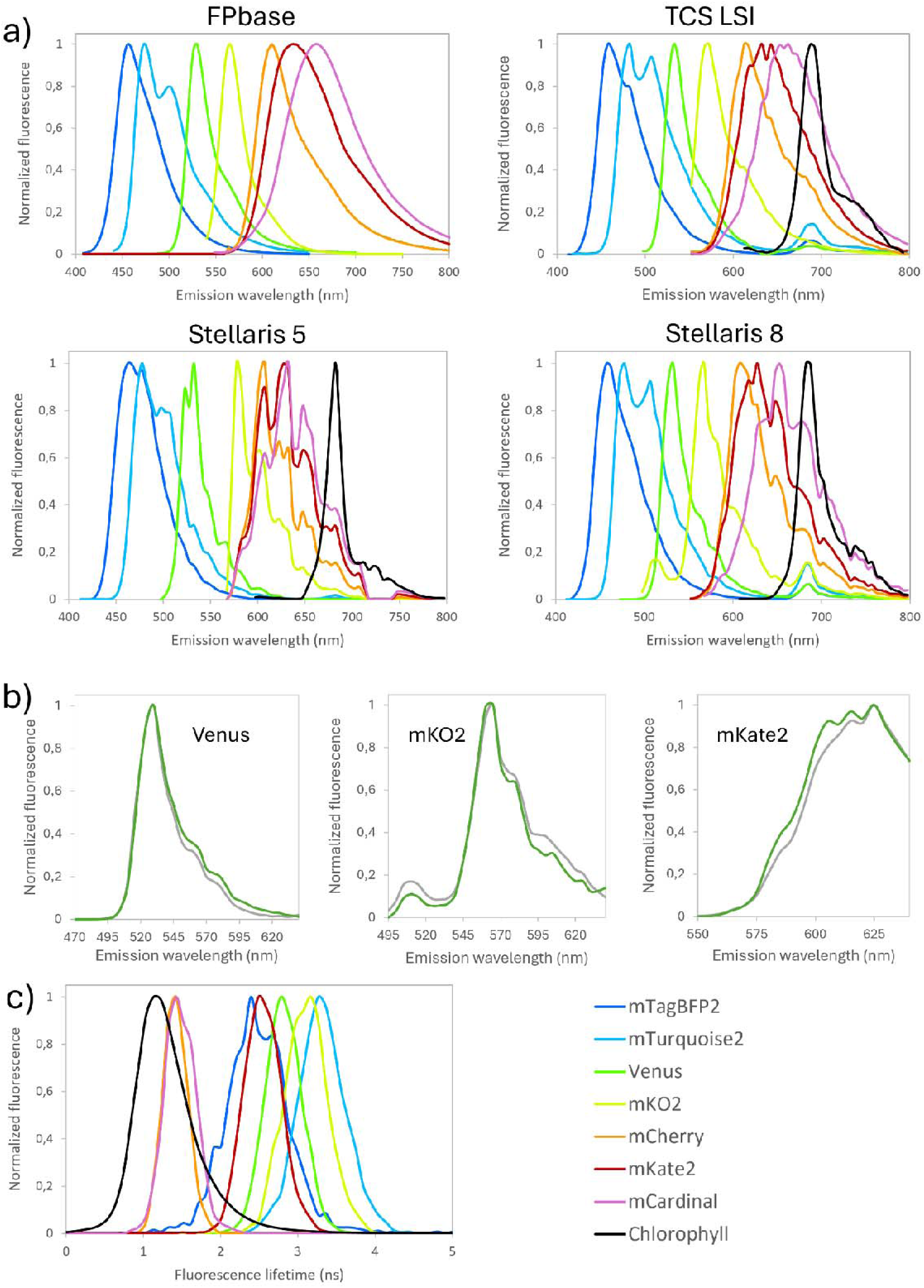
Emission and fluorescence lifetime spectra of a diverse set of fluorescent proteins across confocal systems and plant species. (a) Emission spectra of transiently expressed fluorescent proteins in *N. benthamiana*, measured with optimized settings on Leica’s TCS LSI, Stellaris 5, and Stellaris 8, show imaging system-dependent variations, particularly in the red part of emission spectra compared to FPbase reference spectra. (b) Comparison of three selected fluorescent proteins emission spectra in *N. benthamiana* (green) and potato (*Solanum tuberosum*, gray) reveals similar spectral profiles. (c) Fluorescence lifetime of selected fluorescent proteins and chlorophyll, recorded on Stellaris 8 using white light laser excitation, highlighting clear separation of mKate2 signal from other red spectra emission fluorescent proteins.

To evaluate the influence of the plant matrix, we compared the emission spectra of Venus, mKO2, and mKate2 in stable potato transformants, which closely matched those obtained in transient *N. benthamiana* expression (Figure 1b). These results suggest that while the recorded emission spectra highly depend on the used confocal system, they can be used interchangeably across different plant matrices.

In addition to spectral properties, fluorescence lifetime offers an alternative means of separation. Chlorophyll autofluorescence generally exhibits shorter lifetimes than fluorescent proteins, enabling separation via Tau gating. Our measurements showed that all proteins except mCardinal and mCherry could be fully separated from chlorophyll autofluorescence using Tau gating, even in cases of overlapping emission spectra, such as with mKate2 (Figure 1c). Tau gating imaging can thus aid separation when these proteins are used as reporters.

### All selected fluorescent proteins, except mTurquoise2, show comparable brightness using standard lasers in confocal system

Fluorescent protein brightness is determined not only by extinction coefficient and quantum yield, but also by level of expression, maturation efficiency and pH stability (Cranfill et al., 2016), with additional contributions from laser power and detector sensitivity. Maintaining comparable brightness among spectrally overlapping proteins is essential for accurate separation and broad quantification range: the dimmer fluorophore must remain detectable, while the brighter should avoid oversaturation.

To assess fluorescent proteins’ brightness, we measured cumulative brightness using standard lasers setting (Stellaris 5) or a white light laser or 405 nm laser (Stellaris 8). We limited emission detection to 640 nm to minimize chlorophyll interference. All fluorescent proteins exhibited similar brightness except mTurquoise2 if excited with 405 nm laser, which showed lower brightness compared to FPbase predictions (Figure 2a, b). However, excitation with white light laser (440 nm) resulted in comparable brightness of mTurquoise2 with other fluorescent proteins, although it was still much lower compared to FPbase predictions (Figure 2c, e). These results suggest that although valuable, FPbase predictions might not necessarily reflect the achievable brightness *in planta*.

**Figure 2.**
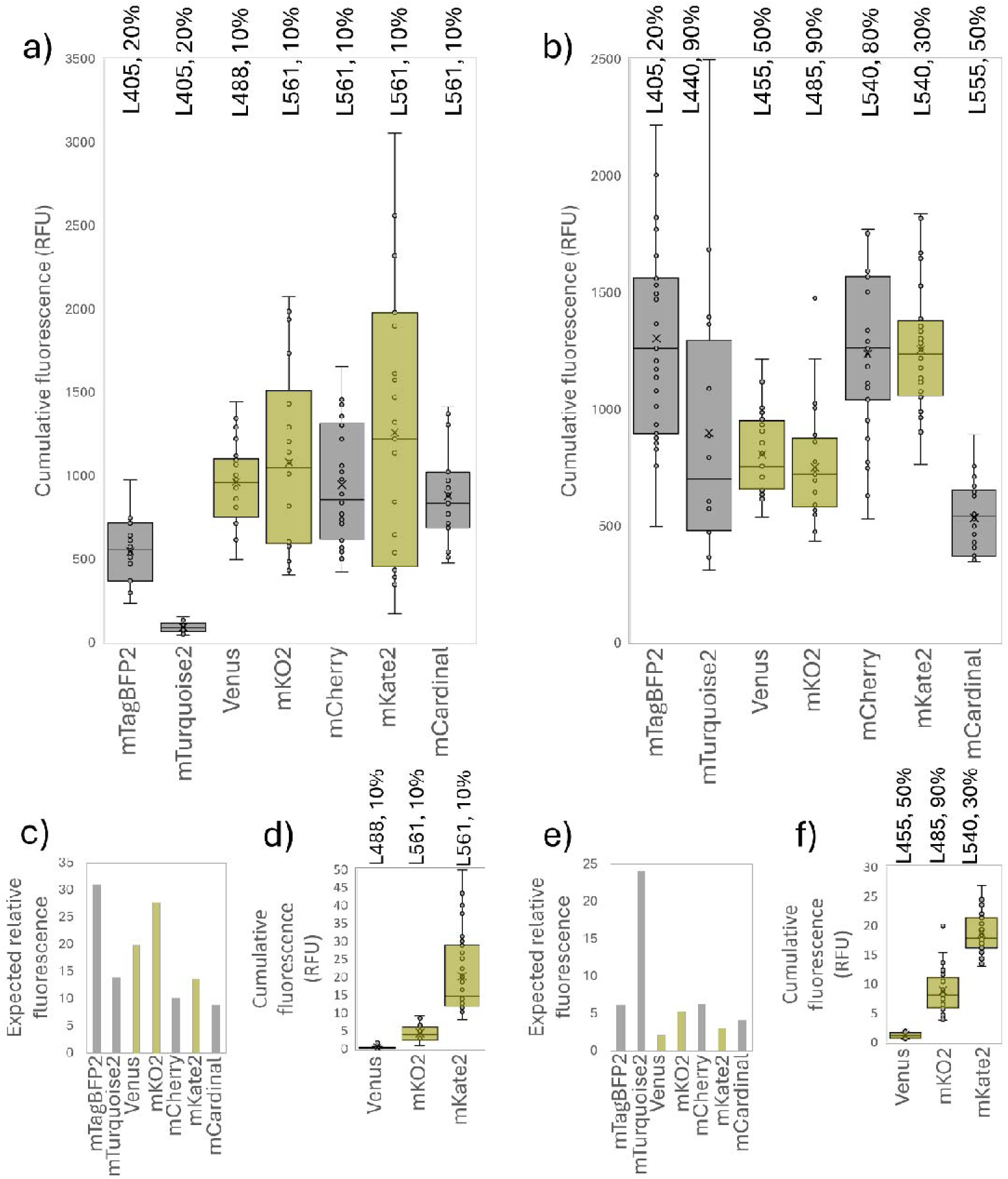
Brightness comparison of fluorescent proteins in *N. benthamiana* and *S. tuberosum* leaves. *mTagBFP2, mTurquoise2, Venus, mKO2, mCherry, mKate2,* and *mCardinal* exhibit approximately comparable brightness across different excitation conditions. All fluorescent proteins were fused to a nuclear localisation signal. (a) Cumulative fluorescence intensity (RFU) of transiently expressed proteins in *N. benthamiana,* measured using standard laser lines on the Leica Stellaris 5 microscope. (b) The same measurement performed using the 405 nm or a white light laser on Stellaris 8, with excitation wavelengths set just below each protein’s emission peak. Fluorescent proteins are indicated below each bar chart, while laser wavelengths and power settings are shown above. Detector gain was kept at the minimum setting for all fluorescent proteins. For each condition, at least two independent images were analysed. Mean fluorescence intensity per nucleus was plotted as bar charts with overlaid points (each dot represents one nucleus). (c, e) Expected fluorescence intensity based on FPbase.org data (excitation × brightness), shown for (c) Stellaris 5 and (e) Stellaris 8, calculated for (a) and (d) standard laser lines, and (b) and (f) white light laser excitation. Among the tested fluorescent proteins, *Venus, mKO2* and *mKate2* were selected for stable transformation in potato (highlighted in yellow across all charts). (d, f) Brightness of fluorescent proteins measured in *S. tuberosum* (potato) leaves (d) using standard laser lines or (f) the white light laser, as described in panels (aLJb). Brightness is shown for one transgenic line per fluorescent protein. Analysis of a second transgenic line gave similar results (results available on Zenodo: 10.5281/zenodo.19691651).

We selected Venus, mKO2 and mKate2 for additional analysis as their brightness was the highest in *N. benthamiana* leaves (Figure 2a; highlighted in yellow in Figure 2). Stable expression of Venus, mKO2, and mKate2 in potato produced fluorescence signals in leaves, which were 100–500× weaker than in transient expression in *N. benthamiana* (see y-axis values in Figure 2d, f versus Figure 2a, b), possibly due to silencing of the transgene (Rajeevkumar et al., 2015).

### Spectrally overlapping fluorescent proteins can be distinguished using linear unmixing

Next, we evaluated the available linear unmixing approaches for separation of fluorescent proteins. Linear unmixing is a powerful method for resolving mixed spectral contributions within individual pixels (Zimmermann, 2005). Previous studies have applied this approach to separate two fluorescent proteins in plant leaves using Olympus and Zeiss systems (Mylle et al., 2013). These authors reported acquiring lambda scans of a combination of two selected fluorescent proteins followed by linear unmixing and its qualitative evaluation. Here, we acquired images by Leica’s Stellaris 5 and applied two linear unmixing strategies. The first one was spectral unmixing, which processes xyλ(z) (lambda scan) images across the entire emission range with fixed wavelength steps. This method works by taking emission spectra (a lambda scan) of every single pixel and then mathematically calculating how much of each known fluorophore (e.g., 60% GFP, 40% Venus) is in each pixel. However, this process is time-consuming, as it requires the microscope to scan the entire emission spectrum for every pixel, typically in 5 or 10 nm steps. The second strategy was channel separation, which processes images obtained by standard acquisition in channels with defined, typically wider emission windows (xy(z) images), where the number of the detection channels must be equal to or greater than the number of fluorophores. This simpler and faster method uses standard channel images (e.g., a ‘green’ channel and an ‘orange’ channel) and calculates a correction matrix based on how much ‘green’ signal is known to bleed into the ‘orange’ channel (and vice-versa) to remove the crosstalk. To set up optimal image acquisition and analysis protocol, we co-expressed three fluorescent proteins: EGFP, Venus, and mKO2, selected for their strong spectral overlap and high brightness. Each protein was targeted to its own cellular compartment; EGFP was localized in the cytoplasm, Venus in chloroplasts (pt-Venus) and mKO2 in nuclei (mKO2-N7). We opted for nuclei and chloroplasts localisation to enable automated segmentation and thus robust quantification. We have previously developed a script for chloroplast segmentation and quantification (Lukan et al., 2023), which we have now adapted for nuclei segmentation and quantification, allowing parallel analysis of up to four different fluorescent proteins (https://github.com/NIB-SI/Nuclei-segmentation). Cytoplasmic fluorescent proteins can alternatively be quantified using mean or cumulative fluorescence across the entire field of view or the remaining part that was not segmented, which is also an available option in the MATLAB script. Apart from co-expression of these three fluorescent proteins, we transformed plants with each of the proteins separately for generation of experimentally determined reference spectra (for spectral unmixing) or reference channels (for channel separation).

We first tested the spectral unmixing approach. As the selected fluorescent proteins have emission peaks close together, the narrowest possible (5 nm) λ-steps for the Leica’s system, were required in image acquisition. Reference spectra were derived from FPbase or acquired experimentally from single-object regions of interest (ROIs) in plants expressing individual fluorescent proteins (Figure 3). Figure 3a illustrates spectral overlap between fluorescent proteins in selected xyλ images: at 510 and 525 nm EGFP (cytoplasm) overlaps with Venus (chloroplasts) and mKO2 (nuclei); and at 560 nm Venus and mKO2 overlap. Using FPbase spectra as references (Figure 3b) resulted in incomplete separation, with residual signals in unintended compartments (arrows, Figure 3b). Improved separation was achieved using experimentally acquired reference spectra (Figure 3c). However, we still observed an insignificant crosstalk signal (Figure 3c, arrows). This is likely due to organelle movement during prolonged imaging, especially at higher magnifications, and reduced signal-to-noise ratio due to narrow λ-steps imaging (Zimmermann, 2005).

**Figure 3:**
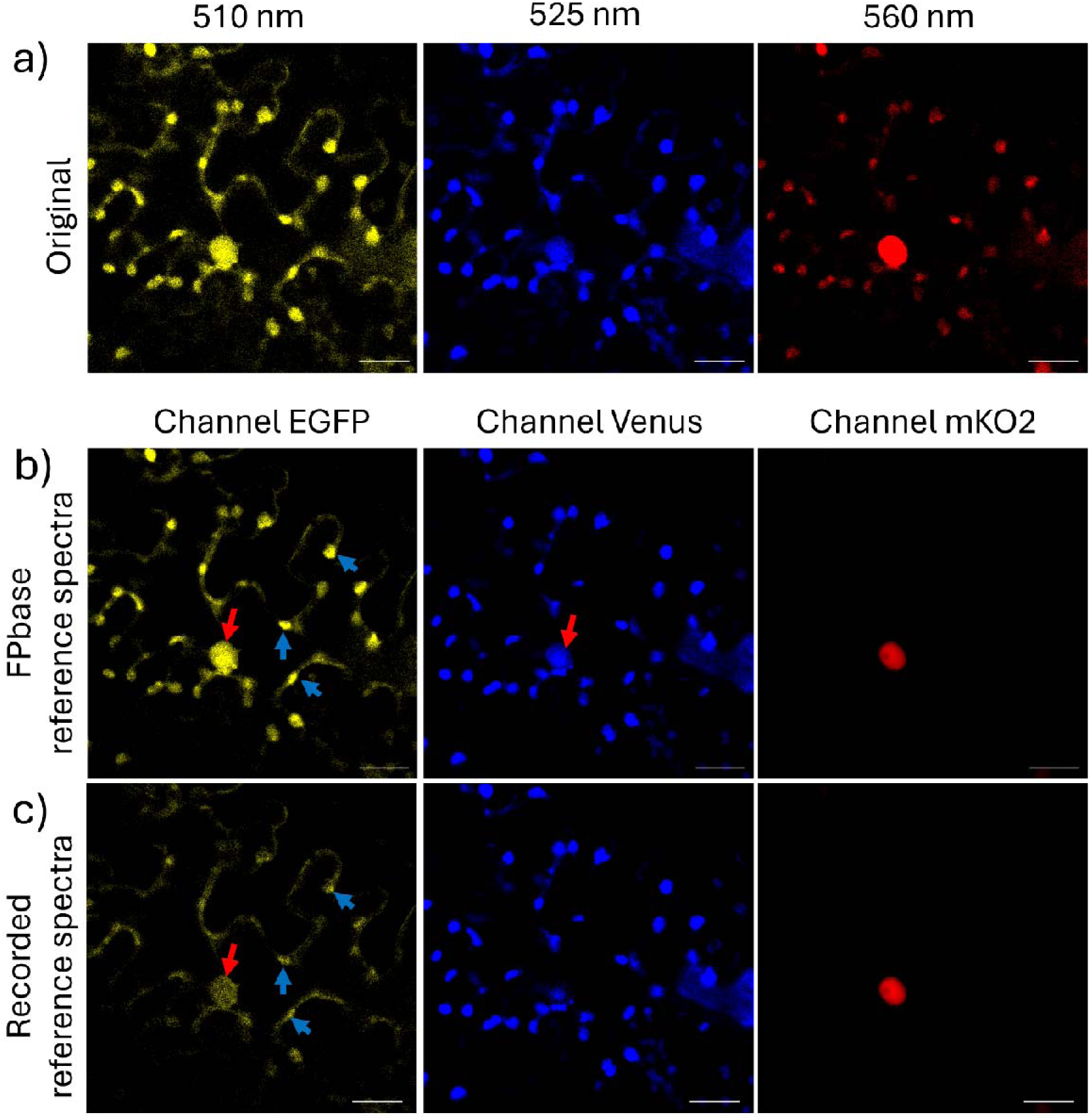
Accurate reference spectra are critical for effective spectral unmixing. (a) Selected images of an xyλ image show crosstalk between fluorescent proteins localized in different cellular compartments. From left to right: images taken at emission peak of EGFP (510 nm), Venus (525 nm) and mKO2 (560 nm). Spectral unmixing results using different reference spectra: (b) FPbase obtained reference spectra, and (c) experimentally recorded reference spectra. From left to right: images obtained after spectral unmixing showing EGFP, Venus and mKO2 channel. Examples of remaining crosstalk from mKO2 in the nucleus and Venus in the plastids after unmixing are pointed with red and blue arrows, respectively. Scale bar: 25 µm.

For unmixing of images obtained by standard acquisition channels (channel separation), reference channels were recorded from plants expressing single fluorescent proteins: (a) EGFP, (b) Venus, and (c) mKO2 (Figure 4). Experimental images were acquired from plants co-expressing all three proteins with the same acquisition settings as for reference channels. In all cases, imaging was performed in three channels with optimized emission windows for the selected fluorophores. Reference channels were then used to manually construct the crosstalk matrix, which was subsequently applied to the experimental images. This approach, which requires substantially shorter acquisition times and can also include channels excited with different lasers, produced results comparable to spectral unmixing using system-acquired reference spectra (Figure 4d-e vs. Figure 3). Quantitative analysis of residual crosstalk error across all unmixing approaches supported the qualitative observations (See Methods for quantification approach; results are presented in Table S1). The most effective separation across all acquired images was achieved using time-demanding spectral unmixing, followed by channel separation, both using experimentally obtained reference spectra. However, even when approximate spectra from FPbase were used, unmixing improved fluorophore separation compared to unprocessed images. For multiplexed fluorescent protein imaging, we therefore recommend channel separation as the preferred method, as its shorter acquisition time reduces motion artifacts and photobleaching while introducing only minimal residual crosstalk. While we used organelle-targeted fluorescent proteins for a robust quantification of unmixing accuracy, linear unmixing can also separate more than three overlapping and colocalized signals with comparable precision. As a proof of principle, we performed channel separation of five fluorescent proteins and chlorophyll (Figure 5). After channel separation, the remaining crosstalk (normalized to the unmixed signal in the main channel) was insignificant (2% for Venus signal in chlorophyll channel, 1% and 4% for chlorophyll signal in mCherry channel and mCardinal channel, respectively, 1% for mKO2 signal in the channels of chlorophyll, mCherry and mCardinal, 3% for mCherry signal in mCardinal channel and 2% for mCardinal signal in mCherry channel; Figure 5). All the remaining combinations of detected signal in other channels resulted in less than 1% of the remaining crosstalk. This demonstrates a significant improvement in signal separation compared to the crosstalk observed before separation (Table 4). However, any residual crosstalk should be taken into account when quantifying individual signals. In addition, crosstalk can be evaluated on a case-by-case basis by comparing subcellular localization at high magnification: strong pixel-wise spatial correlation between channels suggests persistent crosstalk, whereas distinct localization patterns indicate effective signal separation.

**Figure 4:**
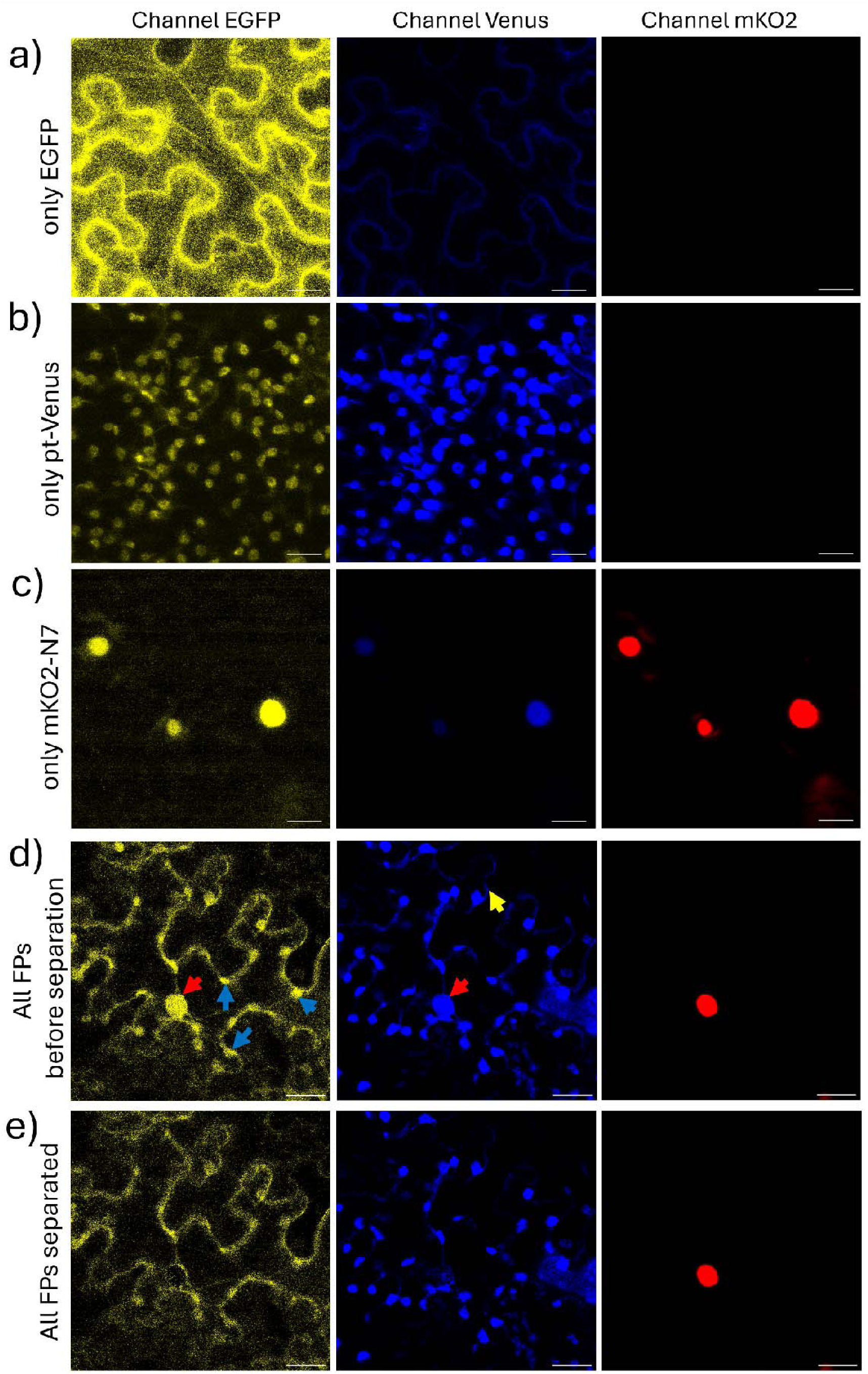
Channel separation produces results comparable to spectral unmixing. (a–c) Reference channels used to generate the unmixing matrix for individual fluorescent proteins: (a) EGFP, (b) pt-Venus, and (c) mKO2-N7. Input (d) and output (e) of channel separation for co-expressed EGFP, pt-Venus, and mKO2. Examples of crosstalk from mKO2 in the nucleus, Venus in the plastids and EGFP in the cytoplasm before unmixing are pointed with red, blue and yellow arrows, respectively. Scale bar: 25 µm.

**Table S1:** Quantification of successfulness of signal separation.

| Original or unmixed set of images | Crosstalk error estimate | Improvement of error compared to original image set (%) |
| --- | --- | --- |
| Original set | 2.29 | 0 |
| Spectral unmixing using FPbase reference spectra | 1.95 | 14.9 |
| Channel separation using recorded reference images | 1.89 | 17.5 |
| Spectral unmixing using recorded reference spectra | 1.80 | 21.2 |

**Figure 5:**
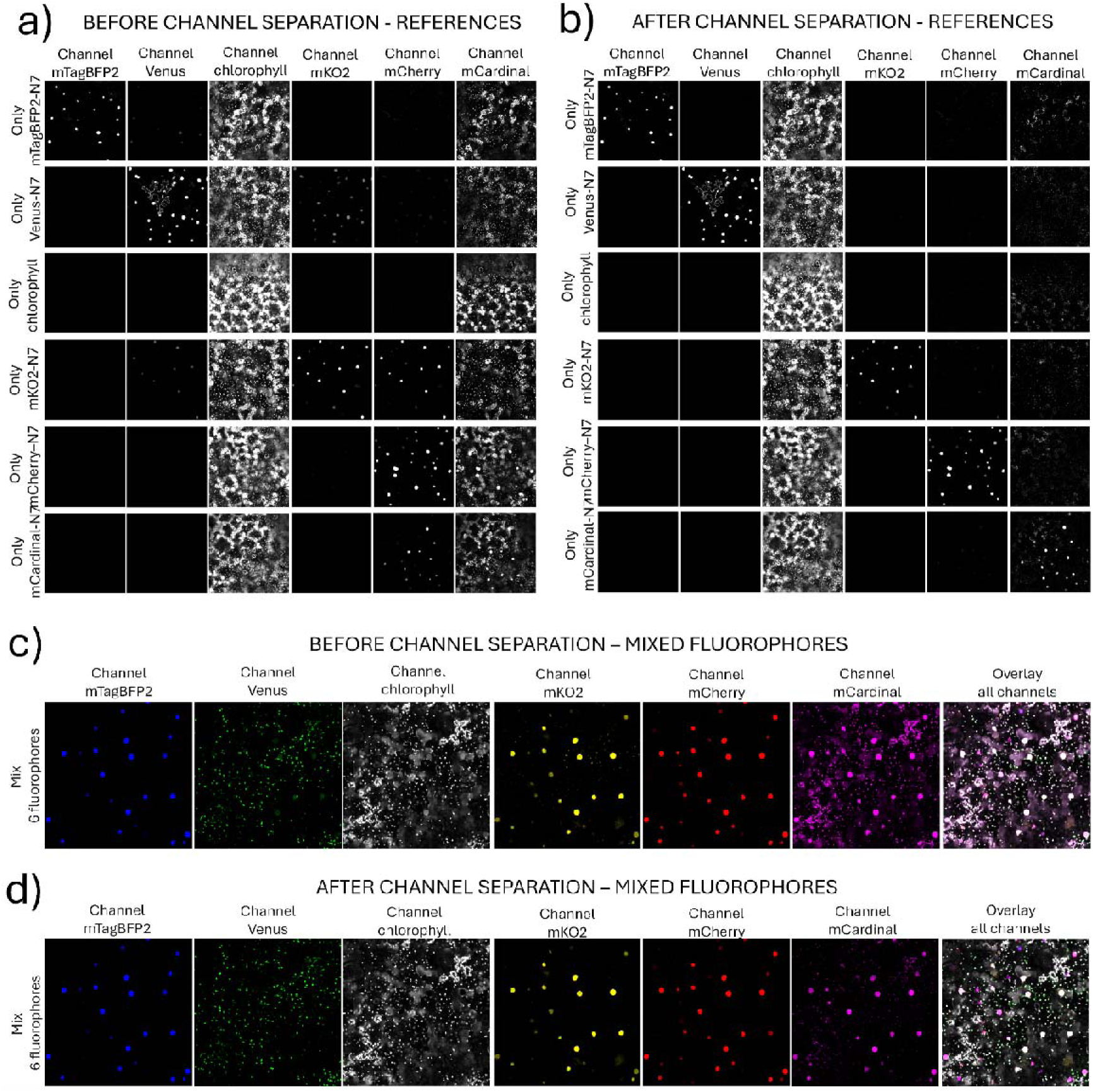
Channel separation enables separation of multiple overlapping fluorescent proteins and chlorophyll. (a) Reference channels of single fluorescent proteins with N7 localization signal for nuclei (mTagBFP2, Venus, mKO2, mCherry, and mCardinal), and chlorophyll channel were used to generate the crosstalk matrix. Images were acquired covering epidermis and mesophyll. (b) Reference channels after channel separation based on the crosstalk matrix produced from images in (a). Note a significant improvement after signal separation. (c-d) Coexpression of five fluorescent proteins (mTagBFP2, mKO2, mCherry and mCardinal in nuclei, and Venus in chloroplasts) and chlorophyll, before (c) and after (d) channel separation with crosstalk matrix calculated from (a). Images were acquired covering epidermis in a single plane. Full image diameter (instead of scale bar): 581 µm.

### Dye separation in practice: monitoring cell dynamics and distinguishing fluorescent proteins from autofluorescence

#### Segmenting nuclei in roots

Finally, we applied linear unmixing under realistic experimental conditions to demonstrate its potential for uncovering new biological insights. As shown earlier, fluorescent protein signals were markedly weaker in potato than in *N. benthamiana*, which limited detection due to a low signal-to-noise ratio. Plant roots are also highly autofluorescent, and in potato, root autofluorescence strongly overlaps with Venus emission, hindering automated nuclear segmentation (Figure 6). Image acquisition on the Stellaris 5 system followed by either spectral unmixing with 5 nm λ-steps or channel separation (Figure 6) increased the contrast between Venus-labelled nuclei and autofluorescent cell walls. Both approaches remarkably improved Venus detectability, enabling reliable automated nuclear segmentation and subsequent quantitative analysis (Figure S1).

**Figure 6:**
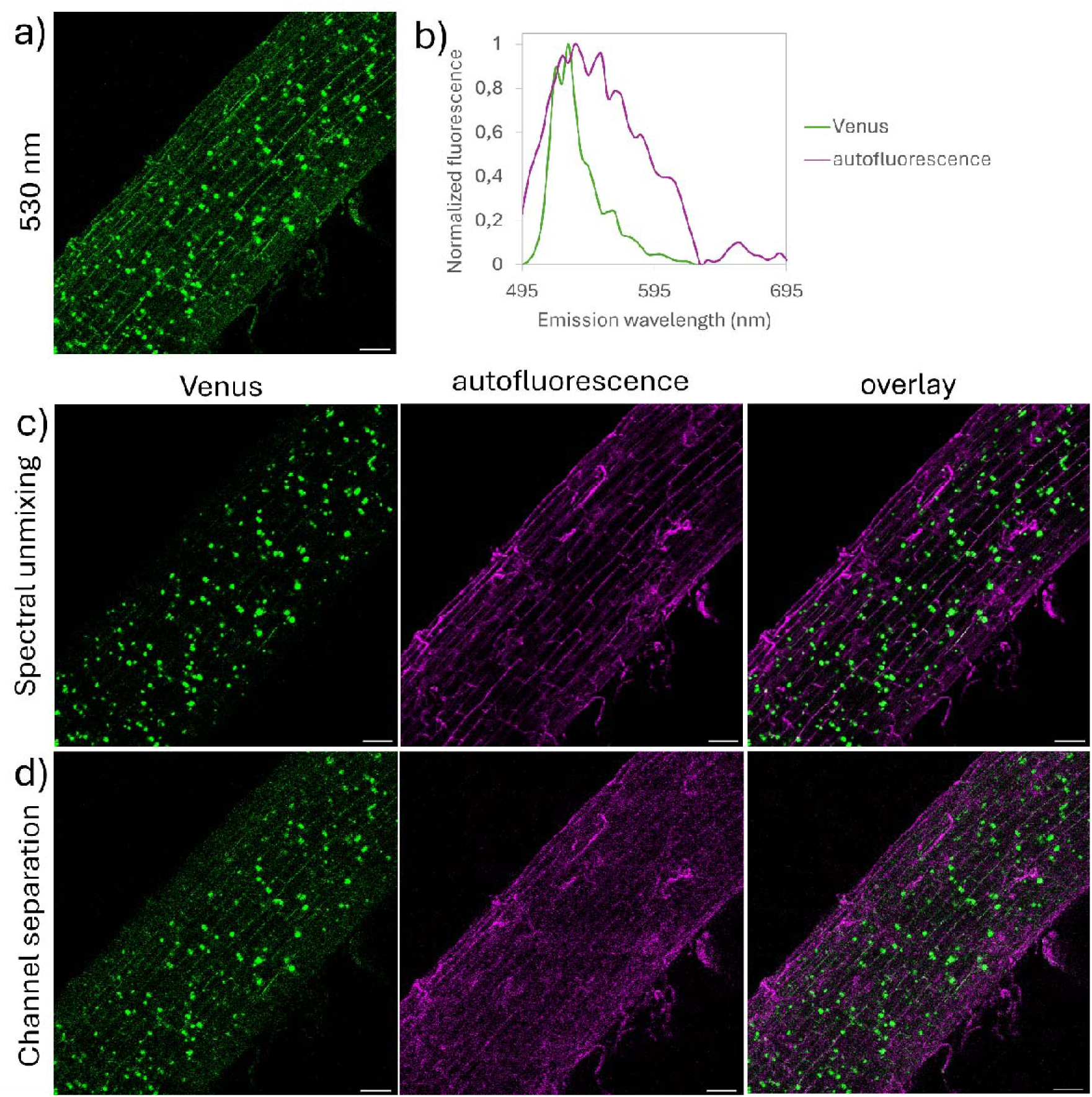
Spectral unmixing and channel separation enhance signal-to-noise ratio for fluorescent protein detection in plant roots. (aDb) Overlap of Venus-labelled nuclear emission (shown in green) with potato root autofluorescence (shown in purple). (c) Spectral unmixing of Venus and autofluorescence. (d) Channel separation of Venus and autofluorescence. From left to right: unmixed Venus, autofluorescence and overlay of both unmixed channels. Scale bar: 100 µm.

**Figure S1:**
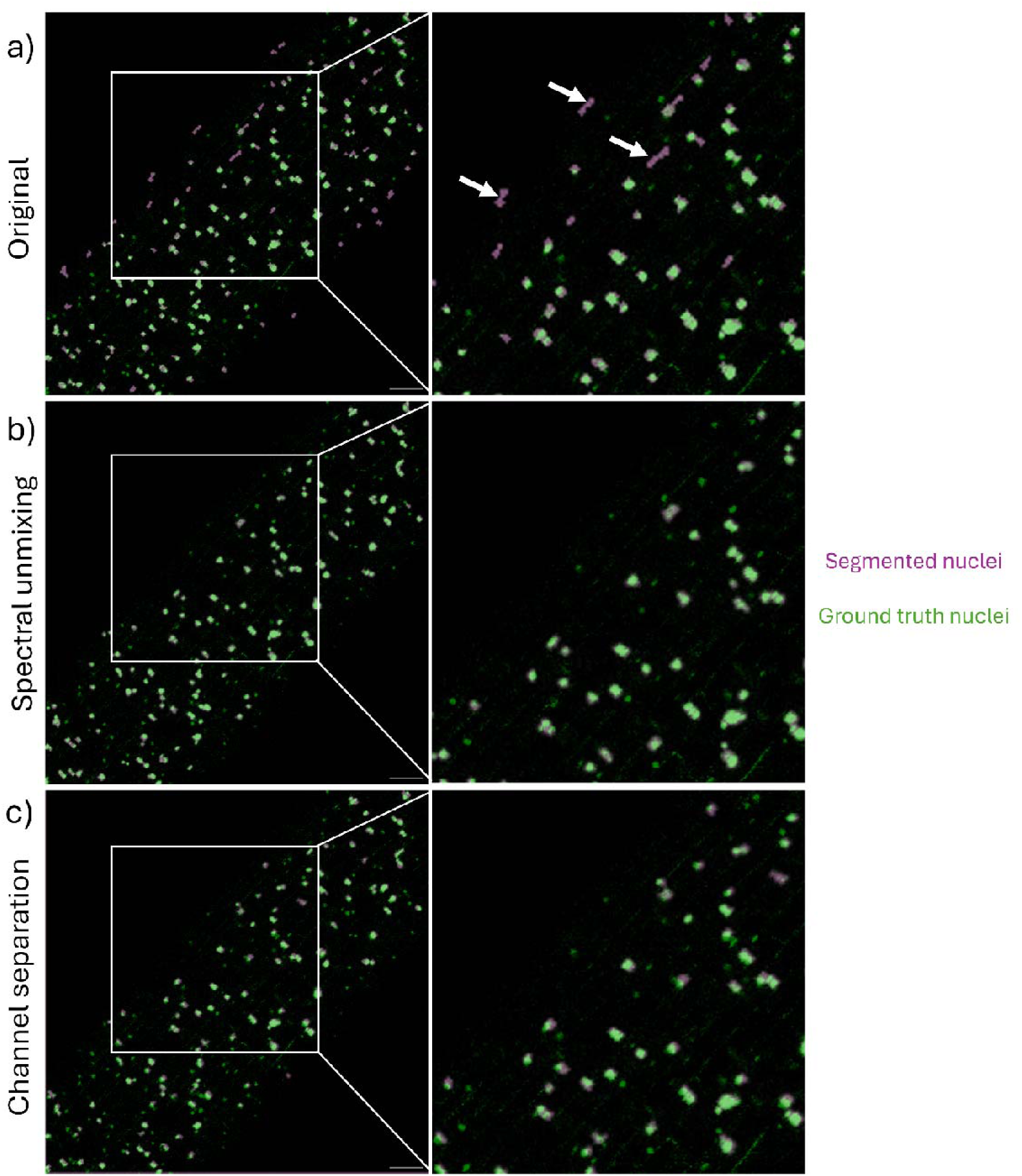
Spectral unmixing and channel separation enable correct segmentation of nuclei in plant root. Ground truth nuclei are marked in green, automatically segmented nuclei in purple. (a) Many nuclei were falsely segmented on image before unmixing (some instances pointed with arrows). Segmentation after (b) spectral unmixing and (c) channel separation is correct. Scale bar: 100 µm.

#### Separation of red fluorescent proteins from chlorophyll autofluorescence

Comparable to nuclear Venus fluorescence from potato root cell walls, nuclear mKate2 fluorescence could be clearly separated from chlorophyll autofluorescence in *N. benthamiana* leaves (Figure 7). In addition to spectral unmixing and channel separation described above, we evaluated a combined approach that exploits fluorescence lifetime differences together with channel separation, using pulsed white-light laser excitation on the Stellaris 8 system. Among the three methods, channel separation yielded the most effective fluorophore distinction (Figure 7). Spectral unmixing performance was limited by sample movement during long acquisition times (arrows in Figure 7 indicate nuclei appearing in the chlorophyll channel). While the Tau gating lifetime-based approach also achieved satisfactory signal separation, it resulted in an overall reduction of fluorescence intensity in both channels, making this combined approach less effective than channel separation alone. Both spectral unmixing and channel separation also enabled complete separation of mCardinal and chlorophyll signals, despite their even larger emission overlap (Figure S2).

**Figure 7:**
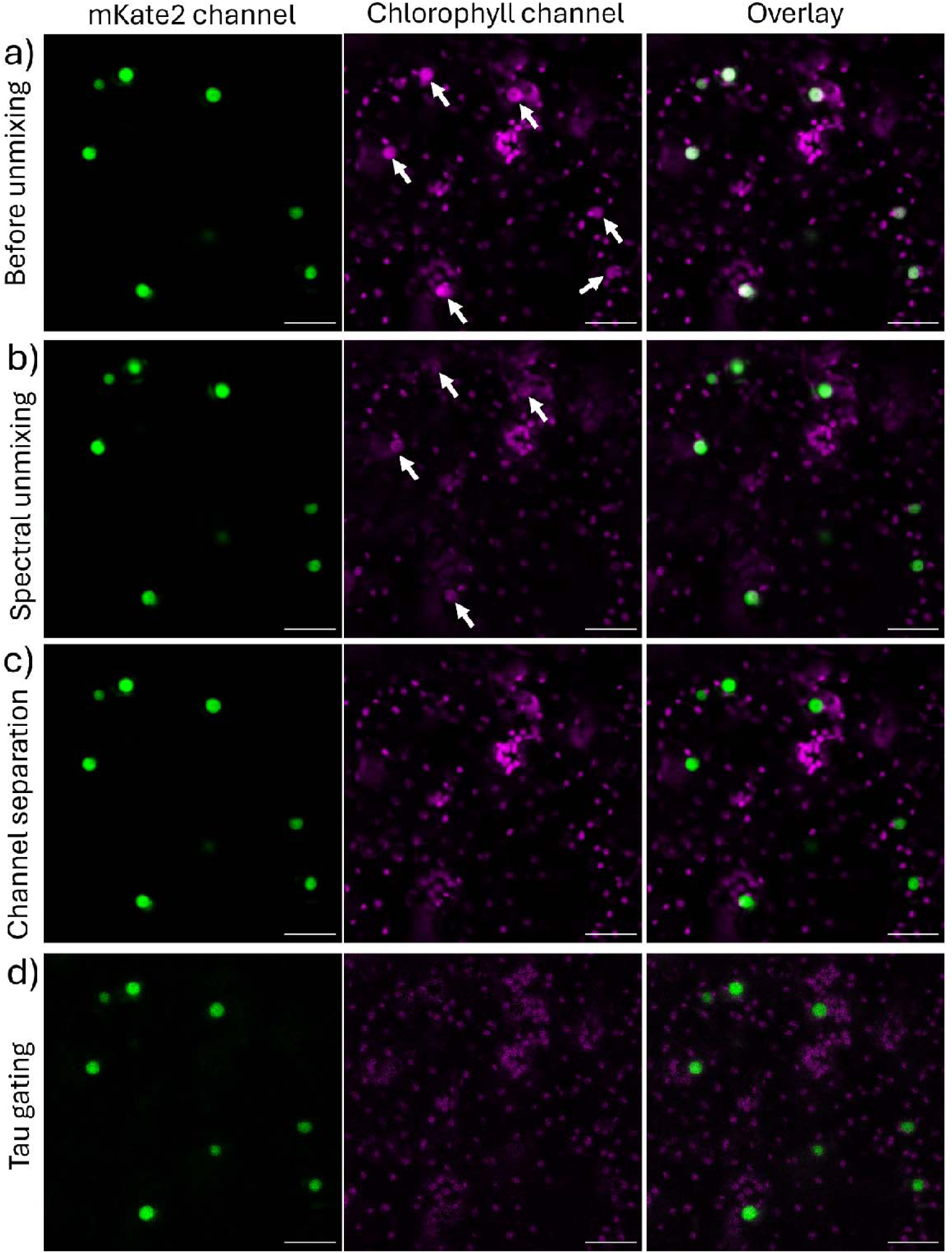
Chlorophyll autofluorescence. 364 **can be unmixed from mKate2 based on spectral or lifetime properties.** Arrows point to mKate2 crosstalk in the chlorophyll channel before and after unmixing. (a) Before unmixing, (b) spectral unmixing using recorded reference spectra, (c) channel separation of mKate2 and chlorophyll, (d) Tau gating followed by channel separation for unmixing. From left to right: mKate2, chlorophyll and overlay of both channels. Scale bar: 50 μm.

**Figure S2:**
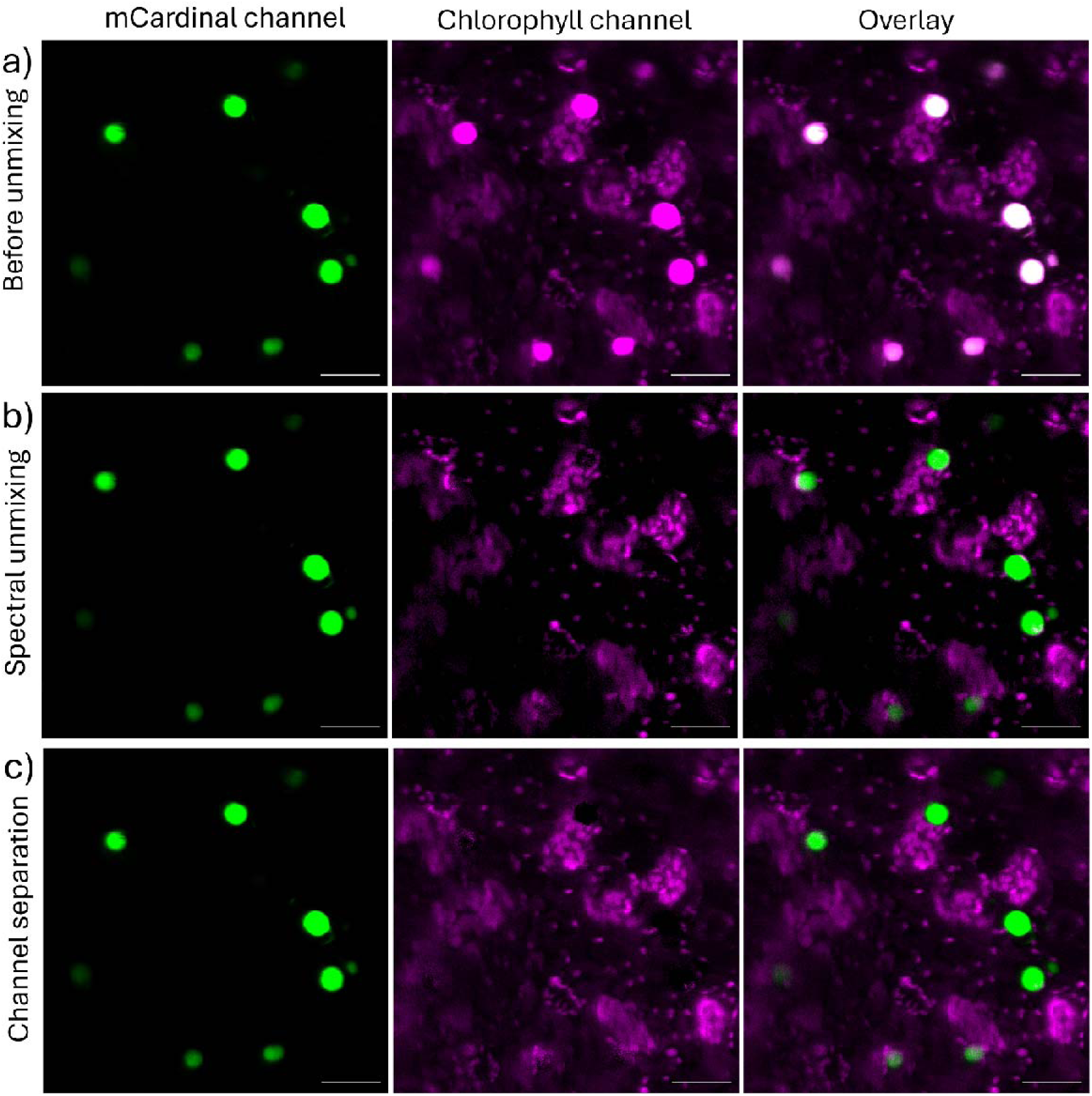
Chlorophyll autofluorescence can be unmixed from mCardinal based on emission spectra. (a) Before unmixing, (b) spectral unmixing using recorded reference spectra, (c) channel separation of mCardinal and chlorophyll. From left to right: mCardinal, chlorophyll and overlay of both channels. Scale bar: 50 µm.

#### Channel separation in live cell imaging

Channel separation can be effectively applied to study organelle dynamics and protein relocalization. As shown in Figures 3b-c and 6b, long acquisitions required for spectral unmixing are prone to disruption by organelle movement and dynamic changes in protein localisation. To overcome these limitations, we recommend simultaneous rather than sequential imaging of overlapping fluorophores, followed by channel separation. This approach enables faster acquisition and thus even real-time tracking of cellular dynamics.

Using co-expression of strongly overlapping fluorescent proteins, pt-Venus and mKO2-N7 located in the plastids and nuclei, respectively, in *N. benthamiana* cells infected with GFP-tagged PVY (Lukan et al., 2023), where GFP localized to the cytoplasm and nuclei of infected cells, we demonstrated that channel separation supports the visualization of rapid and dynamic intracellular processes (Figure 8, Movie 1, Movie S1). Besides faster acquisition, channel separation offers greater flexibility, as it can incorporate fluorescent proteins excited by different laser lines and does not require exact reference spectra, which are strongly system dependent. However, successful application of this method requires reference images of individually expressed fluorescent proteins with comparable brightness in the experimental samples, in order to ensure sufficient signal and avoid signal saturation.

**Figure 8:**
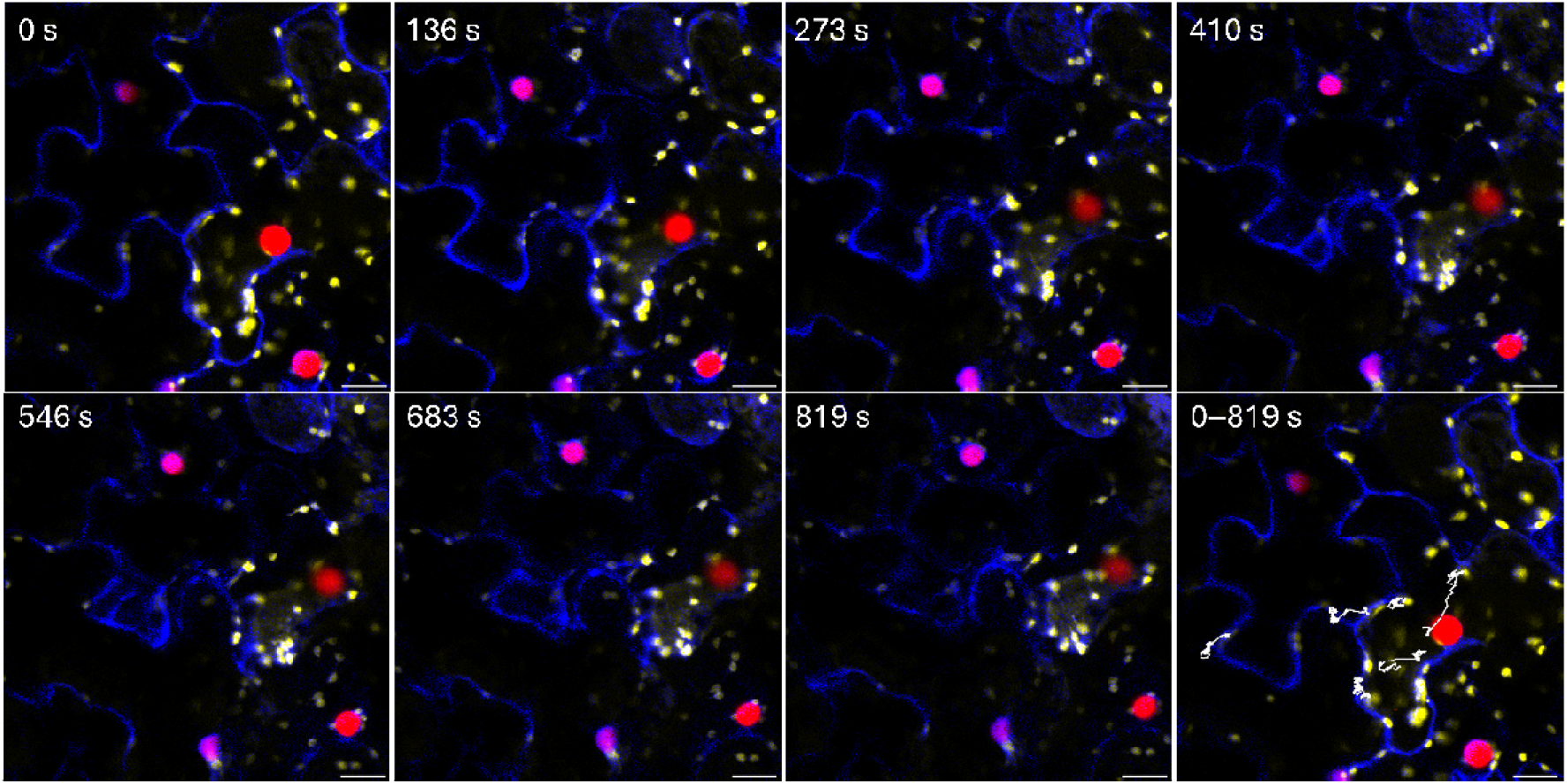
Capturing cell dynamics in PVY-infected cells of *N. benthamiana*. Channel separation enables distinction of cytoplasmic (blue, PVY-GFP) and organellar (yellow, chloroplasts tagged with pt-Venus, and red, nuclei tagged with mKO2-N7) movements in infected cells. Bottom right: tracked movement of selected plastids through time course aligned to the first image (0 s). Scale bar: 25 µm.

## Conclusion

We compared different approaches for separating fluorescent proteins with overlapping excitation and emission spectra in plant leaves. Both spectral unmixing and channel separation effectively distinguished fluorophores, with spectral unmixing achieving the highest accuracy when exact reference spectra were available. However, its application was limited by long acquisition times and motion artifacts, especially at higher magnifications. Channel separation, in contrast, provided comparable separation accuracy with substantially faster acquisition and greater flexibility, allowing the use of multiple laser lines or fluorescence lifetimes. While fluorescence lifetime-based unmixing offered partial improvements, it resulted in lower overall signal intensity.

Overall, our results demonstrate that channel separation offers the best balance between accuracy, speed and robustness and provide a practical framework for optimizing multiplexed fluorescent imaging in plant systems. We showed that five fluorescent proteins and chlorophyll can be separated with insignificant crosstalk. This practical framework would be especially valuable when combining multiple fluorescent protein-based transcriptional reporters in the same plant to follow precise spatiotemporally linked processes, such as different hormonal signalling pathways. Moreover, fluorescent protein multiplexing can prove useful when combining new fluorescent proteins with the existent genetically encoded biosensors which rely on fluorescent proteins with special features (such as fast-maturing Venus in markers of protein degradation (Larrieu et al., 2015; Mir et al., 2016; Song et al., 2021)), e.g. to follow proposed consecutive steps in signalling cascades. However, this approach has limitation for applications requiring high precision, such as colocalization studies, and cannot be used within the same excitation/emission range of ratiometric biosensors such as FRET-based biosensors or roGFP and HyPer and biosensors with normalization channel (Belousov et al., 2006; Hanson et al., 2004; Miyawaki et al., 1997). In such cases, we recommend using non-overlapping fluorophores, such as a combination of mTagBFP2, Venus, and mKate2 or mCardinal, or targeting overlapping fluorophores to different subcellular compartments. To promote reproducibility, we provide MATLAB scripts for nuclear segmentation and signal quantification.

Finally, although databases such as FPbase provide valuable reference data on fluorescent protein brightness and emission spectra, these values may not accurately represent achievable brightness *in planta* or the detected emission profiles. The latter are influenced primarily by the specific confocal system and acquisition settings rather than by the plant tissue itself.

## Materials and methods

### Plant Material and Growth Conditions

*NahG*-Désirée potato plants were obtained from the Leibniz Institute of Plant Biochemistry, Halle, Germany. Désirée, *NahG*-Désirée and newly transformed potato plants were maintained in stem node tissue culture as previously described by Baebler et al. (2009). Two weeks after node segmentation in MS30 agar medium, the plants were transferred to soil and cultivated in a growth chamber under controlled environmental conditions □ 22 °C in the light and 19 °C in the dark, at a relative humidity of 55%, with 120 µmol/m^2^/s radiation for potato (35% white, 5% deep-red and 10% far-red LED lights; PSI, Brno, Czechia) with 16 h photoperiod. *N. benthamiana* were grown from seeds sown to soil and grown as potato plants but with 80 µmol/m^2^/s radiation (20% white, 3% deep-red and 3% far-red LED lights; PSI, Brno, Czechia). Three to five weeks old, soiled potato and *N. benthamiana* plants were used for the experiments.

### Cloning and Vector Construction

Fluorescent protein coding sequences and localization signals were obtained from various sources. N7 localization signal together with mTurquoise2, Venus, and mKate2 (plasmids pHG128, pHG132, and pHG154, respectively) were kindly provided by Hassan Ghareeb (Ghareeb et al., 2016). mTagBFP2 (plasmid pJL1-mTagBFP2; Addgene #102638; (Stark et al., 2018)), mKO2 (plasmid pTU-A-005; Addgene #124412; (Santos-Moreno & Schaerli, 2019)), mCardinal (plasmid mCardinal-pBAD; Addgene #54800; (Chu et al., 2014)), and miRFP713 (plasmid pmiRFP713-N1; Addgene #136559; (Matlashov et al., 2020)) were obtained from Addgene. The plastid transit peptide was amplified from pt-roGFP plasmid kindly provided by Jacob O. Brunkard (Stonebloom et al., 2012), and the H2B nuclear localization signal from the H2B-RFP plasmid kindly provided by Jim Haseloff and Fernán Federici (University of Cambridge, UK).

Fluorescent protein and localization signal coding sequences were assembled through PCR (Table S2) using Phusion® High-Fidelity DNA Polymerase (New England Biolabs, Ipswich, MA, USA) with primers listed in Table S3. PCR products were cloned into the pENTR/D-TOPO vector (Thermo Fisher Scientific, MA, USA) following the manufacturer’s instructions. Constructs were then recombined into binary destination vectors (pB7WG2, pK7WG2, or pH7WG2; VIB Vector Vault, Belgium) via LR reaction (Thermo Fisher Scientific, MA, USA). Chemically competent *E. coli* were transformed with the constructs with heat shock. After growth on selection, plasmids were isolated from transformed *E. coli* colonies using the GenElute™ Plasmid Miniprep Kit (Merck, Vienna, Austria). Sequencing was performed by GATC Biotech using commercial M13 forward and reverse primers (Thermo Fisher Scientific, MA, USA). Finally, constructs were introduced into *Agrobacterium tumefaciens* strains GV3101 and LBA4404 via electroporation for transient or stable transformation of *N. benthamiana* and potato as explained below.

### Transient transformation

For transient transformation, *Agrobacterium tumefaciens* strain GV3101 carrying pB7WG2_Venus-N7, pB7WG2_mTurquoise2-N7, pB7WG2_mCherry-N7, pB7WG2_mKate2-N7, pK7WG2_mKO2-N7, pK7WG2_miRFP713-N7, pK7WG2_mCardinal-N7, pK7WG2_mTagBFP2-N7, pH7WG2_Venus, pH7WG2_pt-Venus, and pH7WG2_H2B-Venus and strain LBA4404 carrying pB7WG2_Venus-N7, pB7WG2_mTurquoise2-N7, pB7WG2_mCherry-N7, pK7WG2_mKate2-N7, pK7WG2_mKO2-N7, pK7WG2_mCardinal-N7, pK7WG2_mTagBFP2-N7, pH7WG2_Venus, pH7WG2_pt-Venus, and pH7WG2_H2B-Venus were grown overnight at 28 °C with shaking at 200 rpm. After 18 h, cultures were centrifuged, and the pellet was resuspended in infiltration buffer containing 10 mM MES, 10 mM MgCl_2_ and 0.15 mM acetosyringone to an OD_600_ 0.3–0.5. For signal comparison, the OD_600_ of Agrobacteria suspensions was adjusted to ensure a maximum difference of 0.05. The suspension was incubated at room temperature with gentle shaking (about 20 rpm) for 2–4 h prior agroinfiltration. Three- to five-week-old *N. benthamiana* were used for transient transformation. For comparison of fluorescent protein brightness only GV3101 strain was used.

### Virus inoculation

On the day of agroinfiltration, four weeks old *N. benthamiana* were first inoculated with PVY-N605(123)-GFP (Lukan et al., 2023) as previously described (Milavec et al., 2008). Briefly, systemic leaves of *N. clevelandii* plants inoculated with PVY-N605(123)-GFP were homogenized in PBS, pH 7.6 (4 ml of buffer per 1 g of infected tissue). The homogenized solution was gently rubbed on the *N. benthamiana* leaf surface sprinkled with carborundum and washed after 10 min.

### Stable transformation

Stable transformations of potato plants cv. Désirée or *NahG*-Désirée with fluorescent protein constructs (pK7WG2_mKate2, pK7WG2_mKO2, and pB7WG2_Venus) were carried out as described in Lukan et al. (2023). Transgenic lines were screened for construct expression using confocal microscopy.

### Confocal microscopy

Confocal imaging was performed using a Leica TCS LSI system with a PMT detector or Leica Stellaris 5 and Stellaris 8 systems with three HyD S detectors. All images were acquired with Plan APO 20×/0.75 NA objectives using LAS X software (Leica Microsystems, Wetzlar, Germany). Exact imaging settings are available in .lif files, image Properties (.lif files are available on Zenodo:10.5281/zenodo.19691651). To enable cross-comparison with other laser scanning confocal systems, we provide the then-current maximum laser powers measured by our service provider: Leica TCS LSI: 405 nm – 2.6 mW, 488 nm – 5.8 mW, 532 nm – 7.0 mW, 635 nm – 2.4 mW. Stellaris 5: 405 nm – 48 mW, 488 nm – 59 mW, 561 nm –165 mW, 638 nm – 159 mW. Stellaris 8: 405 nm – 50 mW, 488 nm – 20 mW, WLL 5 mW. To enable cross-comparison of fluorescent protein excitability primarily based on laser intensity, we performed brightness imaging using 20 nm–wide emission windows and the same detector, whose sensitivity was kept constant across different lasers within the same confocal system.

#### Emission Spectra and Brightness Measurements

Emission spectra were measured in xyλ scanning mode (λ scan) with 5□nm wide steps and 5 nm apart on leaves of transiently transformed *N. benthamiana* plants and stably transformed potato plants. Excitation was performed using 405, 488 or 532 nm lasers (TCS LSI), 405, 488, or 561□nm lasers (Stellaris 5) and 405□nm or a white light laser with wavelength selected just below the emission range (Stellaris 8).

#### Linear unmixing: Spectral unmixing approach

For spectral unmixing, xyλ images were acquired with steps 5□nm wide and 5 nm apart. Experimental and reference spectra were acquired with identical λ-steps to ensure accurate spectral matching. Imaging was performed on Stellaris 5 using 488 nm laser.

#### Linear unmixing: Channel separation approach

For channel separation, images were acquired in multi-channel mode, with each fluorophore captured in its own channel. Detectors gain and emission windows were optimized to balance fluorophore brightness and minimize bleed-through (Table 1, Table 2), ensuring effective separation with minimal signal loss. For separation of mKate2 and chlorophyll based on their fluorescence lifetime, Tau gating was applied at 1.5 ns.

**Table 1:** Optimal confocal settings for Channel separation imaging of 4 fluorophores.

| Fluorophore | Excitation laser (nm) and laser power | Emission window (nm) | Detector gain |
| --- | --- | --- | --- |
| EGFP | 488. 5% | 497–515 | 70 |
| Venus | 488. 5% | 515–547 | 50 |
| mKO2 | 561. 2% | 572–623 | 2 |
| Chlorophyll | 488. 5% | 663–740 | 50 |

**Table 2:** Optimal confocal settings for Channel separation imaging of 6 fluorophores.

| Fluorophore | Excitation laser (nm) and laser power | Emission window (nm) | Detector gain |
| --- | --- | --- | --- |
| mTagBFP2 | 405. 42% | 425–470 | 2 |
| Venus | 488. 2% | 515–520 | 2 |
| Chlorophyll | 488. 2% | 670–730 | 2 |
| mKO2 | WLL 540. 2% | 550–570 | 2 |
| mCherry | WLL 590. 2.5% | 595–635 | 2 |
| mCardinal | WLL 640. 2% | 645–700 | 2 |

WLL, white light laser.

### Data analyses

All results of image analyses have been deposited on Zenodo: 10.5281/zenodo.19691651.

#### Emission and Tau spectra analysis

Average emission and Tau spectra were determined from 15–30 manually selected nuclei across 2–3 micrographs. Chlorophyll autofluorescence was measured in non-transformed *N. benthamiana* by analysing the entire field of view. Average emission and Tau spectra across all nuclei were exported from LAS X (Figure 1).

#### Brightness analysis

Fluorescent protein brightness was quantified as cumulative fluorescence from manually selected nuclei in xyλ images, measured from 5 nm above the laser wavelength (Stellaris 5) or from the beginning of the emission range (Stellaris 8) up to 640 nm to exclude chlorophyll interference (Figure 2). Brightness was determined on 15–30 manually selected nuclei across 2–3 micrographs. Expected relative fluorescence (Figure 2) was calculated from data available at FPbase.org: achievable brightness = brightness × excitation strength at used laser wavelength.

#### Linear unmixing: Spectral unmixing approach

Spectral unmixing was used to separate fluorophores excited by the same laser with overlapping emission spectra. Spectral unmixing was performed on xyλ images using *LAS X Dye Separation* module, *Spectral Unmixing* tool. Reference spectra were obtained from FPbase.org or acquired on plants expressing single fluorescent protein and used by the LAS X Spectral Unmixing tool to mathematically assign emission contributions to individual fluorophores.

#### Linear unmixing: Channel separation approach

Channel separation was used to separate fluorophores with overlapping emission, with excitation from the same or different laser lines using *LAS X Dye Separation, Channel Dye Separation* tool. Channel separation was performed using the *LAS X Dye Separation* module with the *Automatic Dye Separation* tool in *manual* mode. Reference images of single fluorescent proteins were used to calculate a crosstalk matrix (Table 3), with the following assignments: Dye 1 – EGFP (cytoplasm), Dye 2 – Venus (plastids), Dye 3 – chlorophyll, and Dye 4 – mKO2 (nuclei). Channels were defined as Ch1 – primarily EGFP, Ch2 – primarily Venus, Ch3 – primarily chlorophyll, and Ch4 – primarily mKO2. Crosstalk matrix was determined on images acquired on plants expressing single fluorescent protein, normalized to its main channel. More specifically, the brightest object with least chlorophyll bleed-through for each fluorescent protein was manually selected, and its pixel intensities across individual channels were used to build the matrix. Average pixel intensities across individual channels were normalized to the brightest channel of the corresponding fluorescent protein. The matrix was corrected for chlorophyll contributions, which were present in Ch3 for all dyes, by assigning a value of 0 for Dye 1, 2, and 4. This corrected matrix was then applied to unmix images acquired in multichannel mode.

**Table 3:** Matrix used for unmixing of 4 fluorophores.

|  | Ch1 | Ch2 | Ch3 | Ch4 |
| --- | --- | --- | --- | --- |
| **Dye 1** | 1.00 | 0.16 | 0.00 | 0.00 |
| **Dye 2** | 0.41 | 1.00 | 0.00 | 0.00 |
| **Dye 3** | 0.00 | 0.00 | 1.00 | 0.00 |
| **Dye 4** | 0.34 | 0.17 | 0.00 | 1.00 |

For separation of six fluorophores, new reference and experimental images containing individual fluorophores were acquired and crosstalk matrix (Table 4) was calculated based on the signal in the nuclei (fluorescent proteins with N7 nuclear localization marker) or within a large portion of image (chlorophyll). Dye 1 – mTagBFP2, Dye 2 – Venus, Dye 3 – chlorophyll, Dye 4 – mKO2, Dye 5 – mCherry, Dye 6 – mCardinal; Ch1 – primarily mTagBFP2, Ch2 – primarily Venus, Ch3 – primarily chlorophyll, Ch4 – primarily mKO2, Ch5 – primarily mCherry, Ch6 – primarily mCardinal. The matrix was not corrected for chlorophyll contributions. This crosstalk matrix was applied to unmix images of separate (Figure 5b) and mixed (Figure 5d) fluorophores.

**Table 4:** Matrix used for unmixing of 6 fluorophores.

|  | Ch1 | Ch2 | Ch3 | Ch4 | Ch5 | Ch6 |
| --- | --- | --- | --- | --- | --- | --- |
| **Dye 1** | 1.00 | 0.02 | 0.03 | 0.00 | 0.00 | 0.00 |
| **Dye 2** | 0.00 | 1.00 | 0.13 | 0.08 | 0.01 | 0.02 |
| **Dye 3** | 0.00 | 0.00 | 1.00 | 0.00 | 0.00 | 0.35 |
| **Dye 4** | 0.00 | 0.06 | 0.18 | 1.00 | 0.45 | 0.15 |
| **Dye 5** | 0.00 | 0.00 | 0.17 | 0.01 | 1.00 | 0.56 |
| **Dye 6** | 0.00 | 0.00 | 0.41 | 0.00 | 0.23 | 1.00 |

#### Segmentation followed by estimation of error after linear unmixing

Residual crosstalk after channel separation or spectral unmixing was quantified using histogram areas of segmented objects, with each fluorescent protein localized to a distinct cellular compartment. The segmentation and histograms were acquired using MATLAB script (https://github.com/NIB-SI/Nuclei-segmentation). The parameters used in the script to achieve appropriate segmentation are listed on GitHub, Case 1 (https://github.com/NIB-SI/Nuclei-segmentation).

Crosstalk error was quantified as error = sqrt(((µ_1_-µ_teor1_)/σ_1_)^2^+ (µ_2_-µ_teor2_)/σ_2_)^2^), where µ is an average value of histogram area for unmixed signal of the segmented FP in a channel with the residual crosstalk and σ is standard deviation of histogram area for unmixed signal of the segmented FP in the same channel. E.g. crosstalk error of mKO2 was calculated using the average value of histogram area in segmented nuclei in Venus (µ_1_), and EGFP (µ_2_), channel, and standard deviation in the segmented nuclei in corresponding channels. µ_teor_ equals 0 as theoretical remaining crosstalk after unmixing equals 0. Fifteen images were used for quantification of crosstalk error. Error was estimated for each cellular compartment (containing one fluorescent protein) separately and an average error calculated for all three fluorescent proteins is presented in Table S1.

#### Nuclei segmentation in images with root autofluorescence

Automated segmentation of nuclei in root was performed using the provided MATLAB script using the parameters listed on GitHub, Case 2 (https://github.com/NIB-SI/Nuclei-segmentation).

#### Plastid tracking

For visualization and to distinguish plastid movements from sample drift, we used a MATLAB tracking tool (Track2, custom script). Time-lapse confocal stacks were split into individual frames in FIJI (ImageJ) and imported into Track2. A stable nucleus was tracked first to estimate in-plane stage drift (x–y), which was subtracted from all frames for drift correction. We then tracked multiple plastids. Track coordinates were exported to Excel for plotting, and tracks were overlaid on the first confocal frame for visualization.

## Acknowledgements

We thank Tjaša Mahkovec Povalej for her help with experimental work and Miha Tome for reading and commenting on the draft. Potato plants were obtained from the Leibniz Institute of Plant Biochemistry, Halle, Germany (cv. Désirée). This research was funded by the Slovenian Research and Innovation Agency (research core funding No. P4-0165 and P4-0463, projects J4-1777, J4-60073, J4-70169 and ARIS program for young researchers).

## DATA AVAILABILITY

Raw image data supported with metadata were deposited to Zenodo: 10.5281/zenodo.19691651and can be opened with LAS X available at https://www.leica-microsystems.com/products/microscope-software/p/leica-las-x-ls/downloads/. MATLAB script was deposited to GitHub: https://github.com/NIB-SI/Nuclei-segmentation.

